# A temporal atlas and response to nitrate availability of 3D root system architecture in diverse pennycress (*Thlaspi arvense* L.) accessions

**DOI:** 10.1101/2023.01.14.524046

**Authors:** Marcus Griffiths, Alexander E Liu, Shayla L Gunn, Nida M Mutan, Elisa Y Morales, Christopher N Topp

## Abstract

Roots have a central role in plant resource capture and are the interface between the plant and the soil that affect multiple ecosystem processes. Field pennycress (*Thlaspi arvense* L.) is a diploid annual cover crop species that has potential utility for reducing soil erosion and nutrient losses; and has rich seeds (30-35% oil) amenable to biofuel production and as a protein animal feed. The objective of this research was to (1) precisely characterize root system architecture and development, (2) understand adaptive responses of pennycress roots to nitrate nutrition, (3) and determine genotypic variance available in root development and nitrate plasticity. Using a root imaging and analysis pipeline, 4D pennycress root system architecture was characterized under four nitrate regimes (from zero to high nitrate concentration) across four time points (days 5, 9, 13, and 17 after sowing). Significant nitrate condition response and genotype interactions were identified for many root traits with greatest impact on lateral root traits. In trace nitrate conditions a greater lateral root count, length, interbranch density, and a steeper lateral root angle was observed compared to high nitrate conditions. Genotype-by-nitrate condition interaction was observed for root width, width:depth ratio, mean lateral root length, and lateral root density. These results illustrate root trait variance available in pennycress accessions that could be useful targets for breeding of improved nitrate responsive cover crops for greater productivity, resilience, and ecosystem service.

## 2 INTRODUCTION

Agriculture is a significant emitter of greenhouse gasses contributing 11.2% of all emissions in the US and 74.2% of all nitrous oxide emissions (EPA 2022). Intensification of agricultural practices since the Green Revolution with chemical fertilizers, herbicides, and heavy machinery has accelerated soil erosion, quality loss, and pollution (Kaspar and Singer 2015). Adoption of sustainable agricultural practices are urgently needed to reduce net greenhouse gas emissions by at least 43% by 2030 in order to limit atmospheric warming to 1.5°C (IPCC 2022). In recent years, the benefits of cover cropping have been realized with active root systems maintaining ecological processes year-round (Langdale et al. 1991; Gyssels et al. 2005). Despite cover crops being a proximal and impactful solution to more sustainable agriculture practices, minimal crop selection has occurred for optimizing such species as impacts below ground are challenging to measure.

The root system of a plant is vital for capture of water and nutrients. The spatial and temporal organization of the root system, termed root system architecture, greatly affects plant access to these soil resources. As roots develop, they facilitate the modification of the surrounding soil into a rhizosphere that has distinct hydraulic, microbial, and mechanistic properties which differ from those of bulk soil (York et al. 2016; Helliwell et al. 2017). Root system development is highly influenced by plant resource status and the environment and is the product of evolution and adaptation to local soil environments. Characterization and quantification of root development without disrupting the root system architecture is inherently challenging, especially in numbers large enough to query natural variation across a plant species. Therefore, various high-throughput and non-destructive root phenotyping approaches have been developed for assessment of traits of large genetic populations with clear gel, growth-pouches, and rhizotrons (Topp et al. 2013; Atkinson et al. 2015; Seck, Torkamaneh, and Belzile 2020). Using root phenotyping analysis pipelines to characterize and leverage the genetic diversity available in root system architecture is key for breeding more productive and resilient crops.

Field pennycress (*Thlaspi arvense* L.) is a diploid annual species that has promising application as a cash cover crop, which is a cover crop that can additionally provide a harvestable product. Pennycress is found across a wide array of climatypes with most prevalence in temperate regions. It is winter hardy and is therefore an ideal candidate as a fall planted cover crop between maize-soybean rotations in the US Midwest. The potential utility of pennycress is twofold: as a cover crop providing ecosystem services such as reduction in soil erosion and nutrient losses; and as an additional income source for farmers with production of rich oil seeds amenable as a biofuel (30-35% oil) and as a protein animal feed (Frels et al. 2019). Pennycress is potential model cover crop as it is diploid, has a small genome size (539 Mb), is a close relative to the model plant *Arabidopsis thaliana* L., and has a fast generation time of approximately 2-3 months depending on vernalization requirements (Sedbrook, Phippen, and Marks 2014). Recent work has shown that genome editing techniques could rapidly introduce desirable traits into this otherwise mostly wild species (Chopra, Johnson, et al. 2020).

Genetic bottlenecks are a common circumstance in major crop species such as maize, wheat and rice. For these crops, desirable above-ground agronomic traits and yield are the major selection criteria often under high fertilizer and irrigated water regimes. In turn, root system architecture and performance, especially under low input conditions has been largely ignored (Waines and Ehdaie 2007; Eshel and Beeckman 2013). Efforts to widen the genetic pool in these species are being made to introgress near- and distant-ancestral material back into modern crops (Grewal et al. 2018; Yang et al. 2019). In contrast to the conventional cash crops, pennycress is undomesticated and therefore has a wide genetic pool to potentially select from for traits including those for roots. So far, over 500 natural accessions of pennycress have been collected and sequenced for study in research trials (Frels et al. 2019).

To our knowledge, no study exists on the variation of pennycress root systems among collected accession panels. Here, using a 4D gel-based root imaging and analysis pipeline, we revealed root development of pennycress and plastic response to nitrate nutrition in the seedling stage. Among a panel of 24 pennycress accessions, significant variation in root traits and correlation among traits was observed. A significant interaction between genotype and nitrate levels was also observed with common nitrate treatment responses for root traits.

## 3 MATERIALS AND METHODS

### 3.1 Plant materials

Twenty-four pennycress accessions, three spring ecotypes and 21 winter ecotypes, were used in this study. Latitudes of the original collection locations ranged from 61.6 to −51.73. For the original collection locations, 21 accessions were from North America, one accession from South America, and one accession from Europe. Included in the accessions tested is MN106, a winter North American line, which serves as the reference genome for pennycress. All accessions tested have also now been sequenced (Frels et al. 2019; Chopra, Marks, et al. 2020).

### 3.1 Experiment design and growth conditions

Pennycress seeds were surface sterilized with 70% ethanol, then 5% bleach (v/v) for 12 minutes, before being washed three times with double deionized water (ddH2O). Seeds were then transferred to sterile Whatman™ filter paper (Global Life Sciences Solutions Operations UK Ltd, Sheffield, UK) on plates moistened with 0.2 mM CaSO_4_. The seeds were then stratified at 4°C for 3 days in the dark. Stratified seeds were allowed to germinate in the dark at 21/18°C with a day:night cycle of 18/6 h. After 30 h, uniformly germinated seedlings, with a burst testa and maximum 1 mm emerged radicle, were used for the experiments.

Modified Hoagland’s solution imaging gels solidified with 5% Gelzan™ (Caisson Labs, UT, USA) were prepared for the root imaging experiments. For plant nutrition, nitrate concentration of the gels was varied between zero and 5 mM. The high nitrate solution was composed of (in μM) 500 KH_2_PO_4_, 5,000 KNO_3_, 2,000 CaCl_2_, 1,000 MgSO_4_, 46 H_3_BO_3_, 7 ZnSO_4_.7H_2_O, 9 MnCl_2_.4H_2_O, 0.32 CuSO_4_.5H_2_O, 0.114 (NH_4_)_6_Mo_7_O_24_.4H_2_O, and 150 Fe(III)-EDTA (C_10_H_12_N_2_NaFeO_8_). For a low, trace, and zero nitrate Hoagland’s solution, nitrate levels were modified with (in μM) 1,000, 100, and 0 KNO_3_ and replaced with 4,000, 4,900 and 5,000 KCl respectively. A 2.8 mM MES buffer was added to the Hoagland’s solution and adjusted to pH6 using HCl.

The growth vessels were custom 2L ungraduated glass cylinders to which 1L of gel was added. A single germinated seedling was sown per gel onto the gel surface using sterile forceps. Plants were then transferred to the growth chamber with a day:night cycle of 16/8 h at 21/18 °C at a photon flux density of 300 mmol m^-2^ s^-1^ PAR at canopy height. The plants were terminated once the roots reached the edge of the cylinder or imaging plane of view which was after 17 days. A complete randomized block design was used for all experiments.

Three experiments were conducted: the first was a nitrate response experiment with Spring32; the second experiment was a high and low nitrate experiment with Spring32, MN106 and ISU89; and the third experiment with 24 accessions all grown in high nitrate conditions.

### 3.4 Sample collection and harvest measures

For all experiments, root images were collected of each plant at four time points (days 5, 9, 13, and 17 after sowing). The imaging setup consisted of a computer interfacing with an Allied Vision Manta G-609 machine vision camera (Allied Vision Technologies GmbH, Stadtroda, Germany) with a Nikon 60mm f/2.8D lens (Nikon Inc., Melville, NY, USA) and an electronic turntable. The turntable operated in a water-filled tank to correct for light refraction when imaging the glass cylinders. The glass cylinders were partly submerged above the level of the gel when placed in the center of the turntable. A LED flat panel light was used as a backlight to produce grayscale images of the roots with a black silhouette of roots in the foreground against a white background. Root imaging took approximately 2 min per plant with 72 images collected over a 360-degree rotation. The imaging setup used was based on work by Clark et al. (2011).

After root imaging, roots were severed from the shoots and the shoot rosettes were immediately imaged using a flatbed scanner equipped with a transparency unit (Epson Expression 12000XL, Epson America Inc., CA, USA). Rosette size and leaf counts were determined using a modified PlantCV rosette imaging pipeline (Gehan et al. 2017). After shoot imaging, the root and shoot of each plant was dried separately at 60°C for 3 d for determination of dry biomass. Plant measures and extracted traits are detailed in Table 1.

### 3.4 3D root image processing

Collected gel images were reconstructed into 3D models, skeletonised, and traits extracted using GiARoots and DynamicRoots pipelines (Galkovskyi et al. 2012; Symonova, Topp, and Edelsbrunner 2015). GiARoots was first used to scale, crop, threshold, and binarise the gel images. For all roots in this study the RootWorkPerspective mode was used to downsample and reconstruct 36 of the 72 images into a 3D model with an octree node number of 100,000,000, Rotation Axis of −63, or 0, and a top line added to the top of the model. Root system reconstructions were visually inspected for completeness and artifacts. Seven root traits were then extracted from the 3D models using the GiaRoots3D measure function as detailed in Table 1. A further 64 traits were extracted from the 3D point clouds using DynamicRoots with standard parameters (Table 1).

### 3.5 Statistical analysis

Statistical analyses were conducted using R version 4.2.1 (R Core Team 2022); the statistical analysis R codes including the packages needed are available https://doi.org/10.5281/zenodo.7536940. A total of 44 traits from GiaRoots3D, DynamicRoots, PlantCV, plus derived traits are described in Table 1.

## 4 RESULTS

### 4.1 Root nitrate response

In the first experiment, root growth in varying nitrate concentrations was evaluated across time. Spring32 a spring North American line was used as it had seed readily available and represented a spring ecotype. Plants were grown under high N, low N, trace N, and zero N conditions. Root images were taken every 4 d with significant phenotypic variation observed across time and by N condition (Figure 1, Table 2). Lateral roots started to appear between day 5 and day 9. Primary and lateral root total lengths were similar in length at day 9 (Figure 1A). By day 17, root development slowed in the zero N condition (Figure 1A). For all N conditions, lateral roots were the dominant root class for root length from day 13 (Figure 1A).

**Figure 1.**
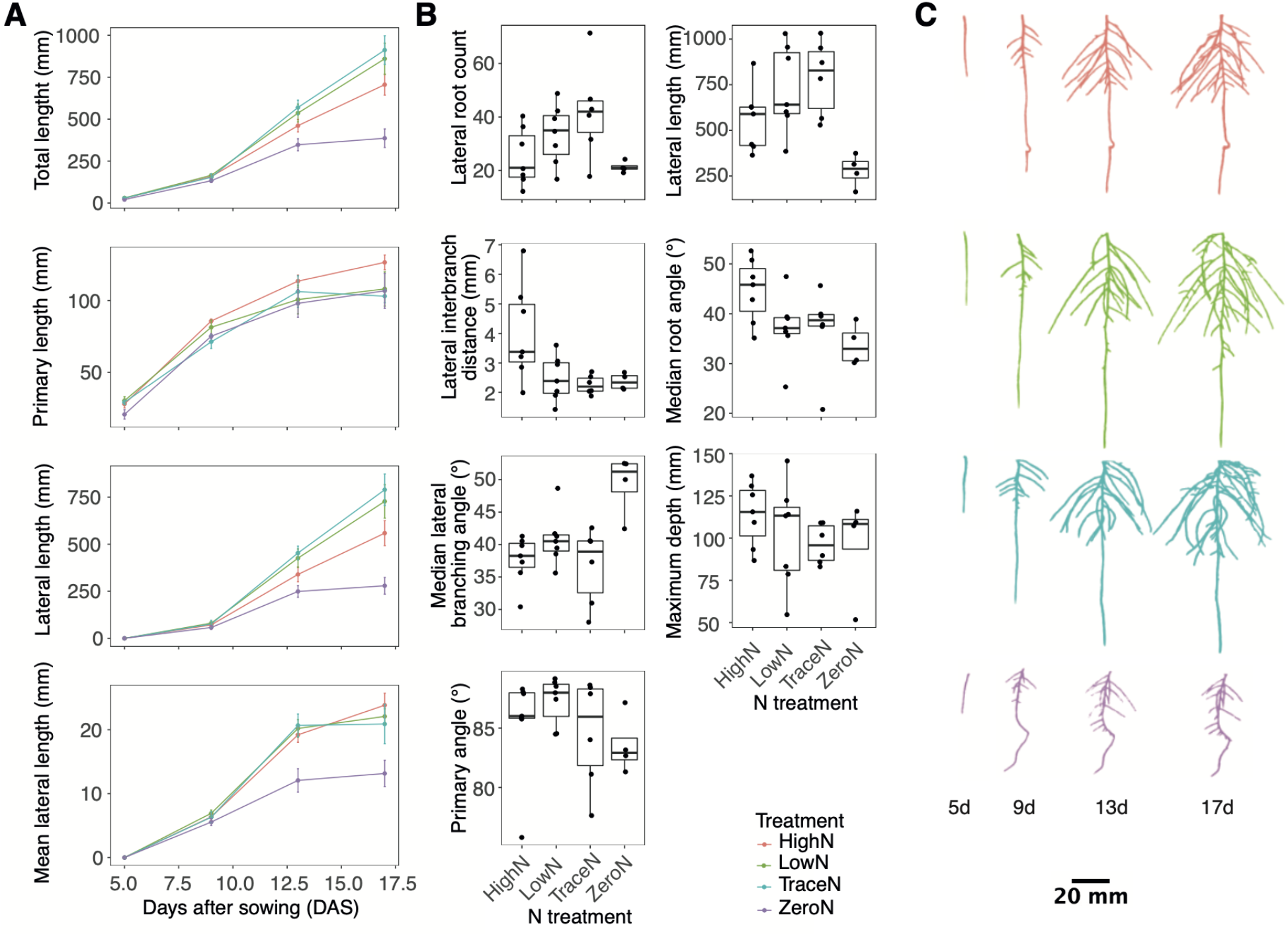
Pennycress root developmental response to varying nitrate concentration. (A) Scatter plots of root traits over time (B) Boxplots of root branching, angle, and distribution traits by varying nitrate concentration at day 17. (C) Representative images of 3D root systems by age and nitrate treatment. High N is depicted as color red, low N as color green, trace N as color blue, and zero N as color purple.

Across the root phenotypes measured, a significant day-by-nitrate condition interaction was observed for lateral root count, length, growth rate, distribution, and angle traits (Figure 1B, Table 2). The greatest lateral root count was observed in the trace N condition with a 51.3% greater count than high N condition and a 65.6% greater count than the zero N condition (Figure 1B). Lateral root interbranch distance was smallest observed in the trace N and zero N conditions with a 56.3% and 51.9% smaller distance respective to the high N conditions (Figure 1B). The greatest lateral root length was observed in the trace N condition with a 34.2% greater length than the high N condition and 95.5% greater length than the zero N condition (Figure 1B). For root angle traits, the zero N condition had the steepest median lateral root angle at 33.7° with a 10.9° difference from the shallowest under high N conditions (Figure 1B). In addition, the zero N condition also had the greatest lateral root branching angle from the parent root with a 11.7° greater branching angle than the high N condition (Figure 1B). For the primary root traits, the differences were less, with the deepest roots observed in the high N condition with a 17% greater root depth than the zero N condition, and the primary root angle was shallowest in the zero N condition with a 3.7° shallower primary root than the low N condition (Figure 1B).

Next, we evaluated GxE responses to nitrate nutrition in the key lines Spring32, MN106, and ISU89, which are important lines as they represent spring, winter, and southern ecotype diversity, respectively. The three accessions were grown under high N and trace nitrate conditions. Root images were taken every 4 d with significant phenotypic variation observed by genotype and nitrate condition (Figure 2, Table 3). All accessions had a greater lateral root length and count under trace N conditions compared to high N conditions (Figure 2A, Table 3). For width:depth ratio a significant genotype-by-nitrate condition interaction was observed (Figure 2A, Table 3). All accessions had a greater width:depth ratio in trace nitrate conditions over high nitrate with ISU89 having the greatest increase of 74.9%. MN106 had the smallest increase in width:depth ratio with a 25.9% increase in low nitrate conditions.

**Figure 2.**
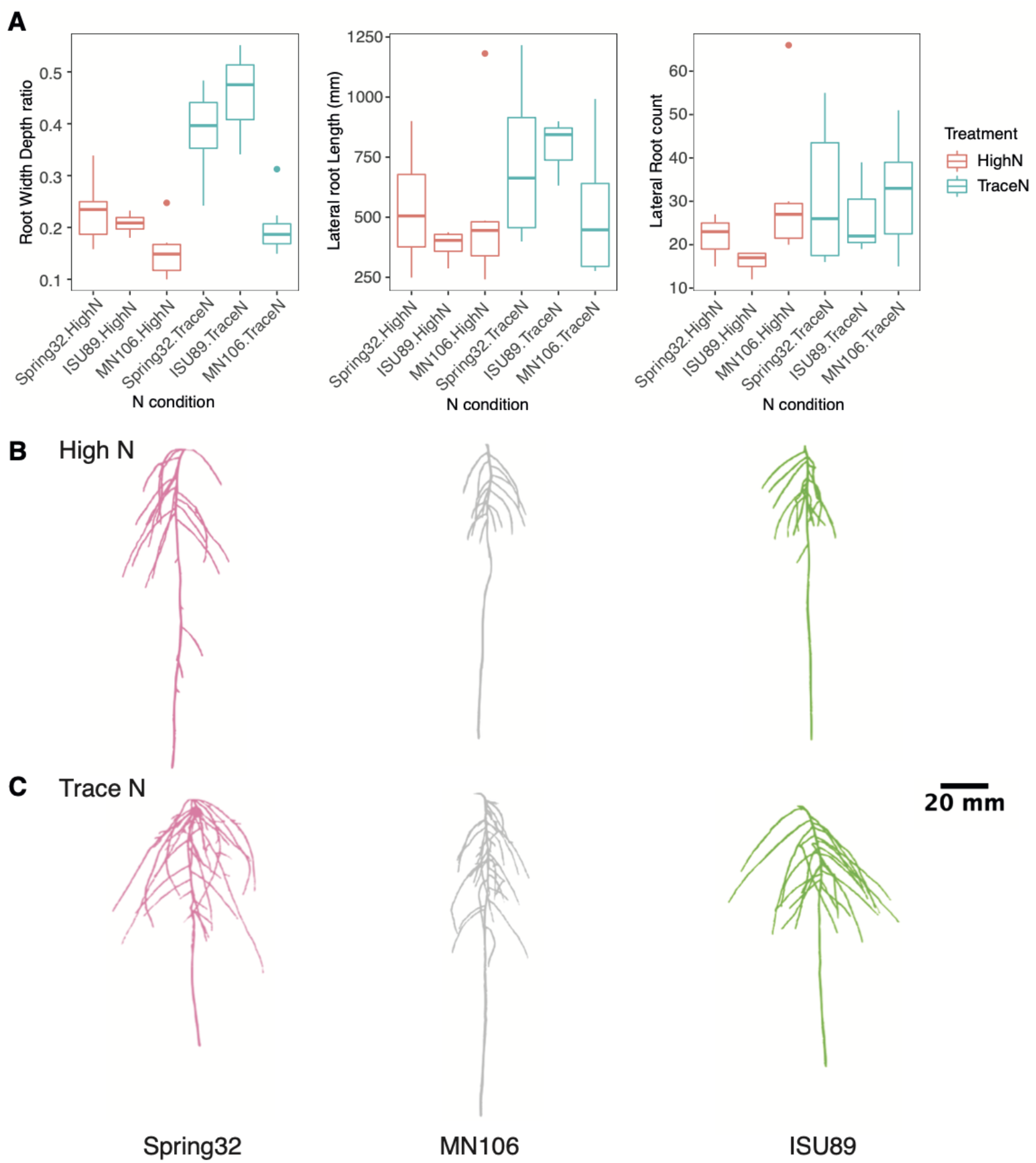
Genotypic and N condition variance in pennycress lateral root traits. (A) Boxplots of accession root traits by nitrate treatment. Representative images of 3D root systems under (B) high and (C) trace nitrate concentrations.

### 4.2 Phenotypic variation among genotypes

In the large experiment, 24 diverse accessions were grown in high N conditions. Root images were taken every 4 d with significant phenotypic variation and visual differences observed by genotype and across time (Figure 3, Figure 4, Table 4). For root size traits including length, radius, and volume; and root distribution traits including bushiness, convex hull volume, width, tortuosity, soil angle, and branching angle; a significant genotype-by-day interaction was observed (Table 4). With a limited number of spring ecotypes tested, separation between spring and winter ecotypes was observed in a linear discriminant analysis with the spring ecotype in general larger than winter types (Figure3ABC). The spring ecotypes had on average a 47.2% greater width:depth ratio, 43% greater lateral root length, and a 25.4% greater root count (Figure 3AB).

**Figure 3.**
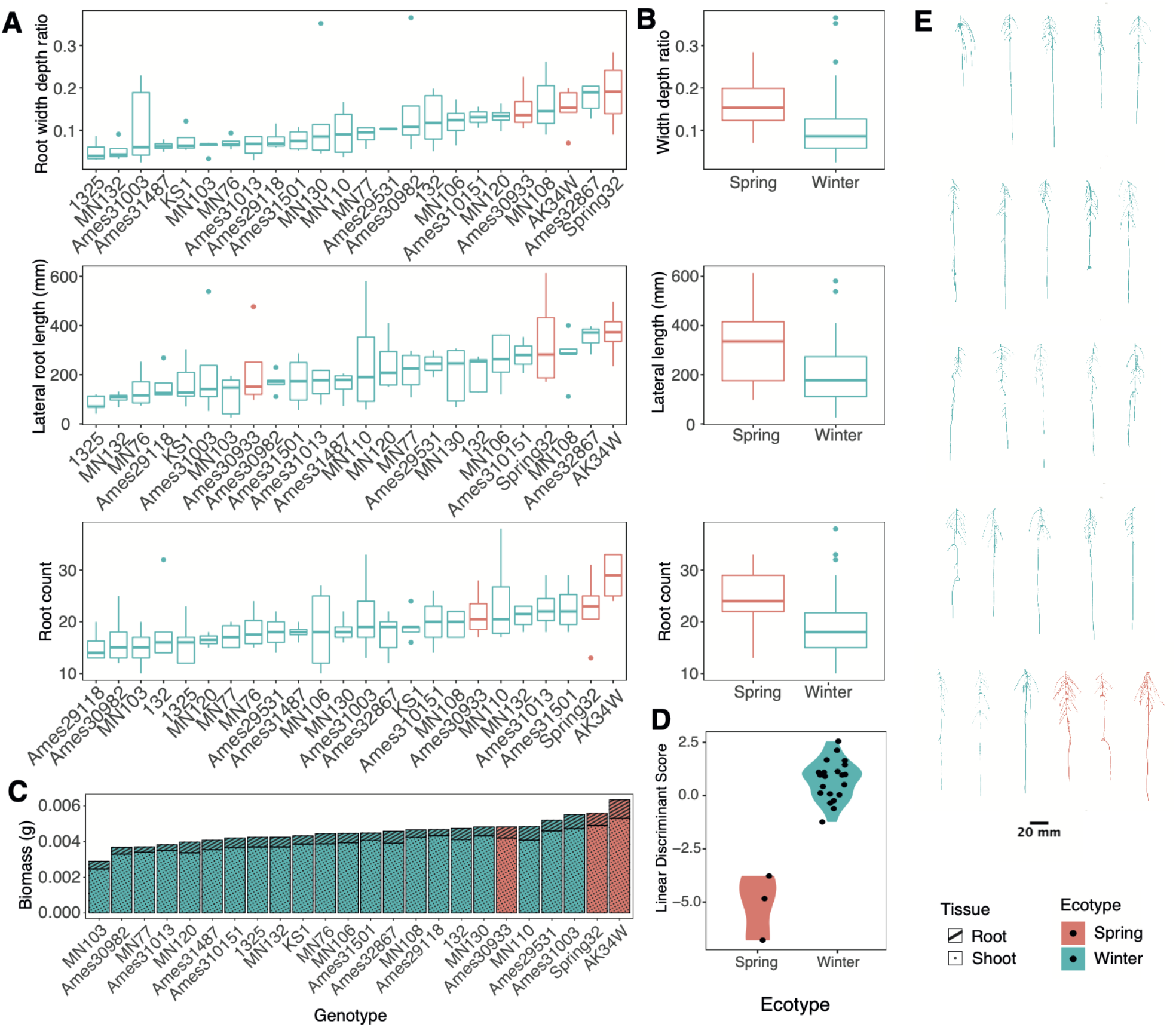
Variance of spring and winter pennycress ecotypes for root traits at 17 days after sowing. (A) Boxplots of root measures of individual accessions. (B) Means of individuals summarized by ecotype. (C) Stacked bar chart of mean root and shoot biomass allocation per plant. (D) Linear discriminant analysis between ecotypes. (E) Representative images of 3D root systems of pennycress accessions. Boxplots and root images are color coded by winter or spring ecotype.

**Figure 4.**
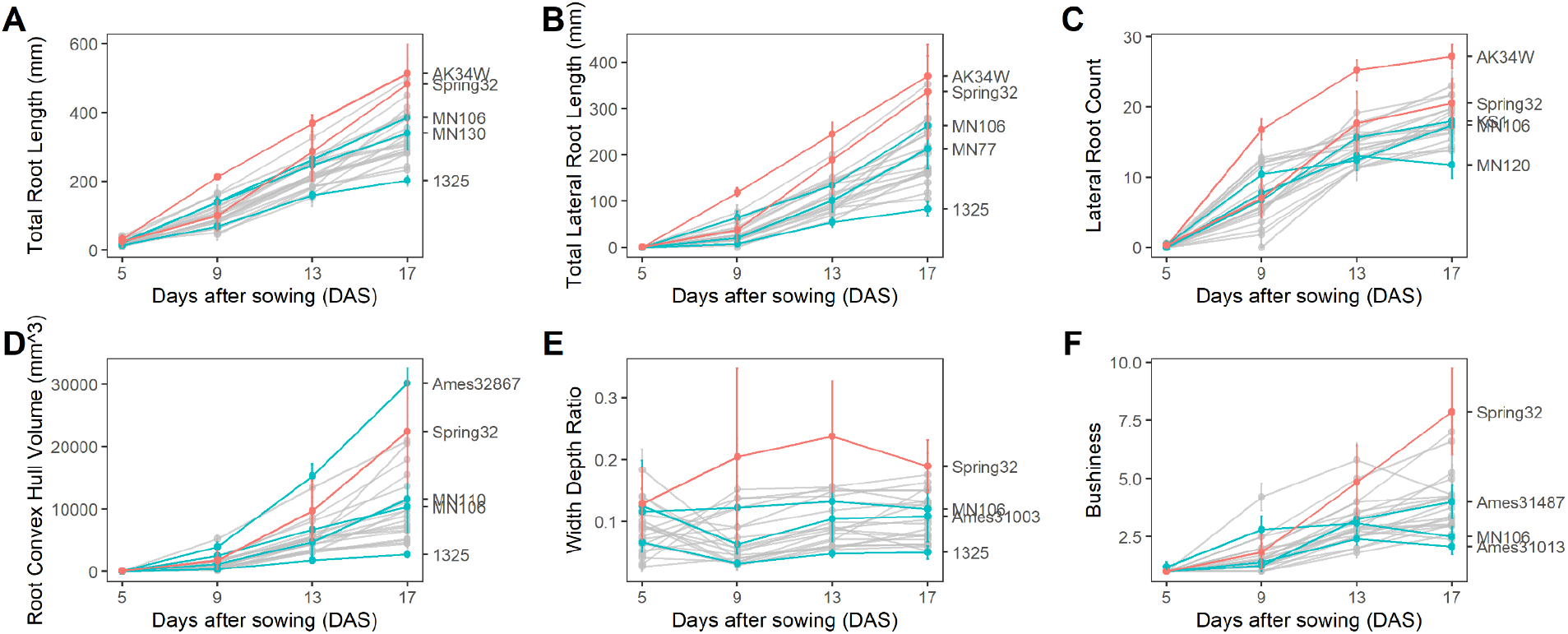
Root phenotypes of diverse pennycress accessions at 5,9, 13, and 17 days after sowing with mean and extreme genotypes highlighted along with Spring32 and MN106 reference lines. (A) Total Root Length. (B) Lateral Root Count. (C) Lateral Root Count. (D) Root Convex Hull. (E) Width Depth Ratio. (F) Bushiness.

By imaging roots over a time course, a diveristy of changes in root traits between genotypes over time was observed. Over time, total root length increased at a constant rate, lateral root production decreased, root convex hull volume increased at an accelerated rate, and width to depth ratio stayed mostly constant. Genotypes with at the extremes of some traits tended to be extreme in other traits. For example, the genotype AK34W demonstrated larger root systems when measured across several traits including total root length, total lateral root length, and lateral root count. Likewise, the genotype 1325 demonstrated a smaller root system based on total root length, total lateral root length, convex hull, and width depth.

### 4.3 Correlation among traits

To determine relationships among pennycress plant traits, correlation and principal component analysis (PCA) were conducted using root phenotyping data of 24 diverse pennycress accession lines. A correlation analysis showed a strong positive correlation among root system size and distribution traits including convex hull volume, root width:depth ratio, and root length (Figure 5A). Root mean radius was positively correlated with total volume, width:depth ratio, and interbranch distance which may be driven by variation in the primary root. Interbranch mean distance was positively correlated with root length distribution. For root depth, a positive correlation was observed between root median lateral root branching angle and a negative correlation with root solidity and root mean radius. For the distribution traits, there was no significant correlation observed between root system width and root system depth (Figure 5B). A positive correlation was observed between root system width and lateral root length (Figure 5C).

**Figure 5.**
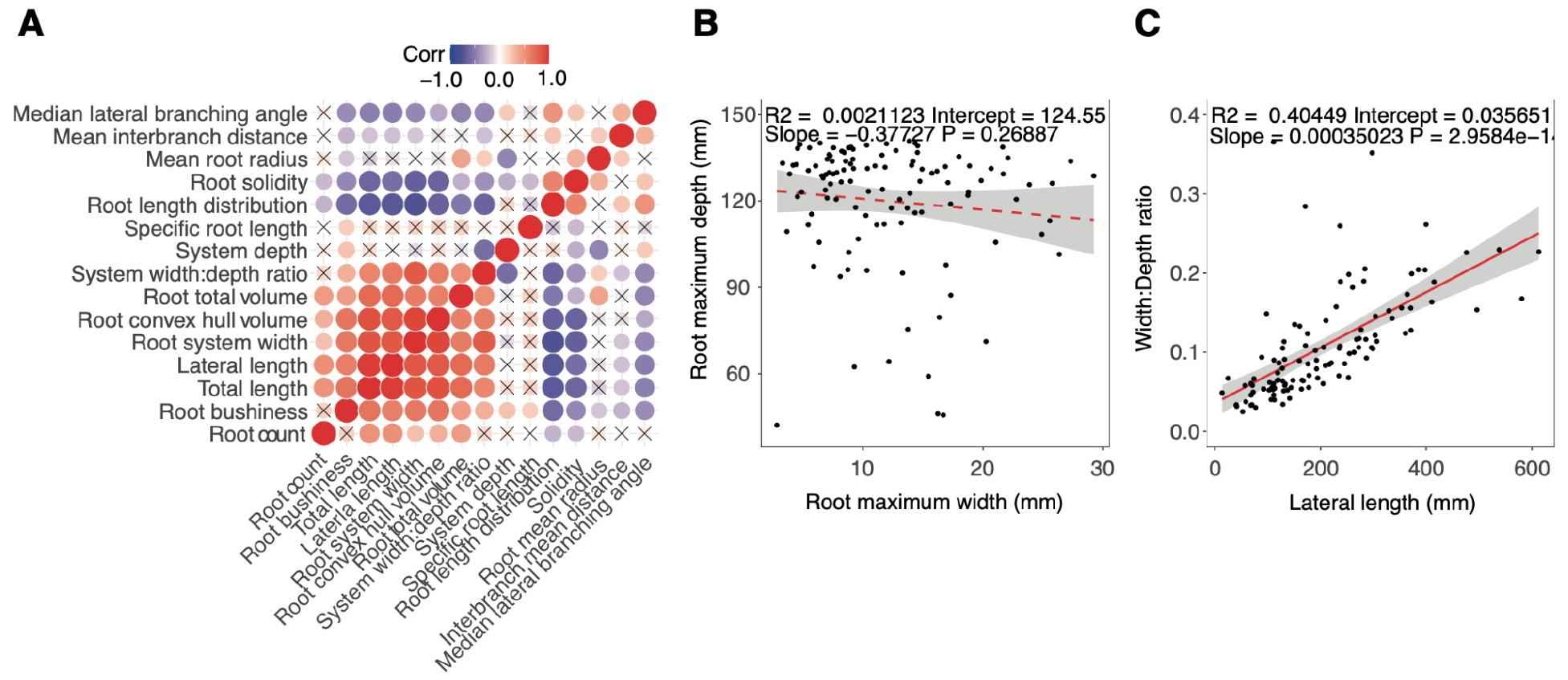
Phenotype correlation between diverse pennycress accessions at 17 days after sowing. (A) Correlation matrix. (B) Regression between maximum root system width and depth. (C) Regression between root system width and lateral root length.

Principal component analysis was used to explore relationships among the plant traits and to explore if there was clustering by pennycress population structure traits (Figure 6AB). With a subset of traits selected to avoid collinearity, the first two principal components explained 60.2% of the trait variation observed (Figure 6A). Over 75% of the trait variation could be explained by the first four principal components. The loadings for PC1 were mostly root size and width traits including root system and lateral root length, root system maximum width, bushiness, and convex hull volume (Figure 6C). Root depth, width:depth ratio, and root mean radius, and solidity were the main traits contributing to PC2 (Figure 6D). A PCA individuals plot was conducted with clustering based on ecotype (Figure 6B). Based on the phenotypes collected at this seedling stage minor separation in clustering was observed between spring or winter ecotypes.

**Figure 6.**
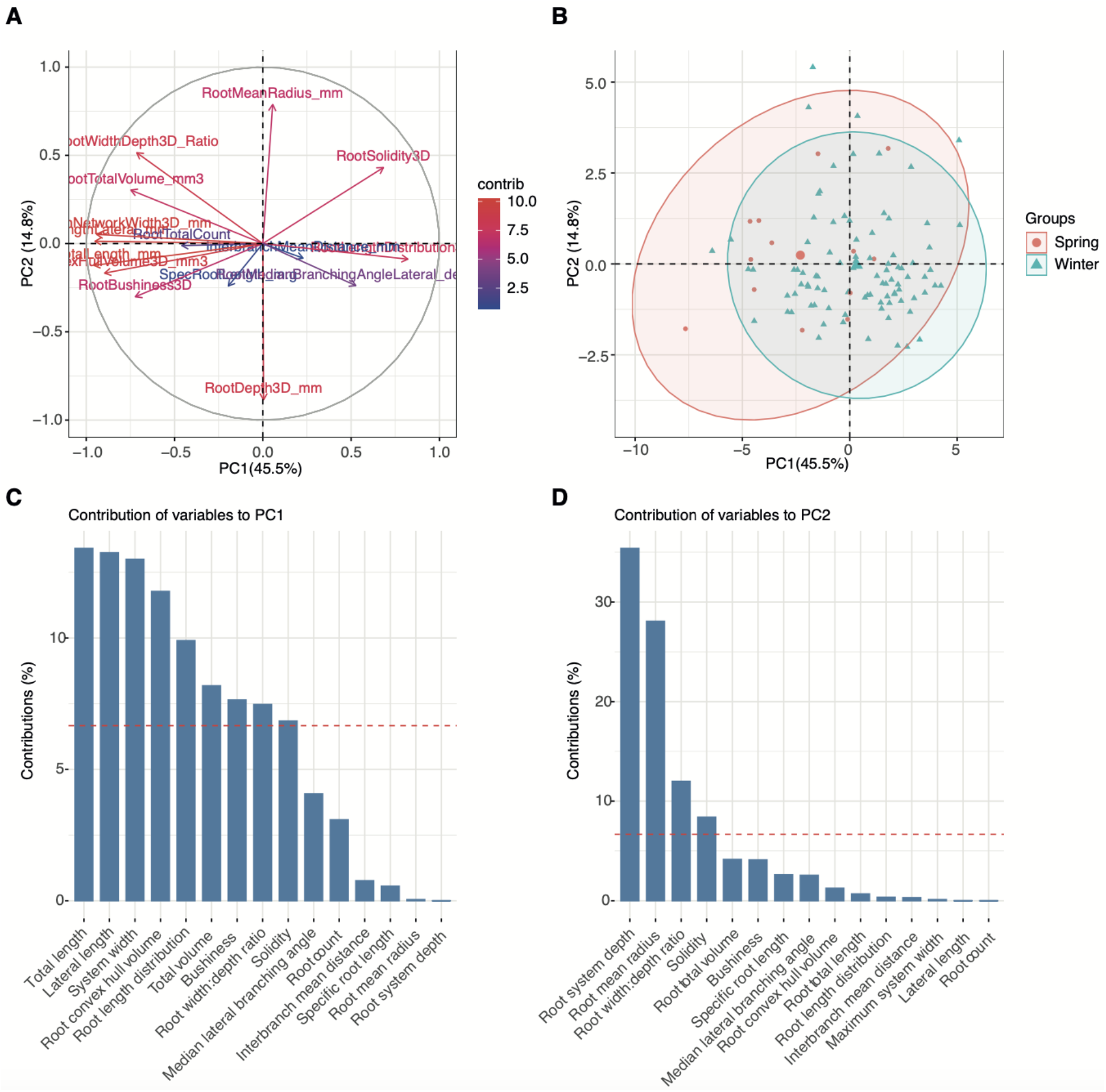
Phenotype association between diverse pennycress accessions at 17 days after sowing. (A) PCA plot of variables. (B) PCA biplot of individuals with ecotype clustering. (C) PCA contribution plot for PC1. (D) PCA contribution plot for PC2.

## 5 DISCUSSION

The spatial and temporal arrangement of root systems are critical for their ability to capture transient and unequally distributed soil resources needed for plant growth and productivity. In turn, root systems are a primary input of soil carbon and drive biological, chemical, and physical changes to the soil which have major ecological implications. Evaluation of in-situ root development at high precision is challenging and requires specialist growth and imaging approaches. Here we used a 4D gel imaging platform to evaluate root system development, architecture, and response to nitrate availability across diverse pennycress accessions.

The primary reasons for adoption of cover crops are to prevent soil erosion and provide other ecosystem services, most of which are directly related to root system architecture and function (Griffiths, Delory, et al. 2022). Yet very little is known about root system development of most cover crops or how they influence the environment. Evaluation of pennycress root traits will provide insights into functional mechanisms and may provide opportunities for improvement of ecosystem services and yield. Pennycress is a tap-rooted plant and here we show that the lateral root system was the dominant root class after 13 days of growth. Pennycress seedlings were grown under varying nitrate concentrations and the lateral root system was most responsive, a phenomenon also observed in Arabidopsis, wheat, barley, maize, rice, and others (Drew and Saker 1975; Trachsel et al. 2013; Yu et al. 2014; Griffiths, Mellor, et al. 2022; De Pessemier et al. 2022). In pennycress, three distinguishable nitrate plasticity responses were observed; (1) normal root development under sufficient or excess available nitrate, (2) intense root foraging to increase access to nitrate for stress avoidance in nitrate limited conditions, and (3) root growth arrest when nitrate became critically insufficient. As nitrogen is heterogeneously distributed in agricultural soil, both spatially and temporally, these plastic responses in root system architecture can facilitate greater uptake with responsive root exploration combined with expression of nitrate transporters (Yu et al. 2014; Melino et al. 2015).

Under trace nitrate conditions an intensive root foraging strategy was observed with greater lateral root count, lateral root length, smaller interbranch distance, and steep lateral root angles. Steeper root angles and biomass allocation has been reported in various studies under nitrogen stress and is positively correlated with nitrate uptake at depth (Trachsel et al. 2013; Griffiths, Wang, et al. 2022). Deep rooting is an ideotype proposed for more nitrate stress tolerant crops and could in turn be used for reducing nitrate runoff as an ecosystem service (Trachsel et al. 2013; Griffiths, Delory, et al. 2022). Root development has a cost however and there must be enough available nitrate for roots to continue development. In the trace nitrate conditions, more nitrate was required for adequate development yet enough was available to facilitate the extra root development. Before day 13, root development was similar across each nitrate condition, however as seen in the zero-nitrate treatment the seed nitrate reserves were likely expired after 13 days as evidenced by root growth arrest. In the high nitrate conditions, a lower lateral root length and interbranch distance was observed compared to the trace nitrate conditions. This indicates that sufficient nitrate was available to the plant and available in the local root zone. Therefore, extra roots were not needed or had diminishing returns with resources likely allocated for other developmental processes.

Pennycress has a wide geographic distribution with collections mainly from temperate regions in the northern hemisphere. The accessions evaluated in this study represent a small proportion of the natural variation available for breeding. Here among a subset of the accessions it was demonstrated that there are common and differential root plastic responses to nitrate concentration. Common responses include changes in root distribution traits including depth, root length distribution, lateral root angle, convex hull, and solidity. For width:depth ratio of the root system a genotype-by-nitrate condition interaction was observed. All accessions tested had a greater width:depth ratio under low nitrate attributed to the greater lateral root length, but the magnitude of change differed between genotypes. These differential root responses to nitrate indicate different tolerance strategies that may in turn affect situational nitrate capture effectiveness or stress tolerance. Harnessing plasticity of yield impacting traits is important for developing generalist genotypes that produce well matched phenotypes to the environment or specialist genotypes that are stable in a trait of interest. Generalist genotypes will outperform specialist genotypes in variable environments and specialist genotypes outperform in more stable environments (Wuest, Peter, and Niklaus 2021; Schneider 2022). Measurement of nitrate uptake performance with isotopes and extending the study to the full lifecycle would provide insight into how these root responses affect plant uptake performance and yield traits.

Comparing root phenotypes of 24 pennycress accessions originally collected from locations in North America, South America, and Europe, significant variance was observed for most measured traits. These differences observed likely represent adaptations to the local environments in which they were collected. Significant root trait differences were observed for most root traits with exception of primary root angle which grows vertically into the ground. Roots have a central role in multiple ecosystem processes and the variance observed in pennycress root traits will likely affect such ecosystem service performance. A correlation matrix and PCA analysis using root phenotypic data from all accessions revealed two root trait spectra for breeding: (1) root size, density, and width traits and (2) root depth and lateral root angle traits. Denser rooting is one such trait that has desirable application for sustainable agriculture by reducing soil erosion rate (Langdale et al. 1991; Gyssels et al. 2005). Deeper and more extensive rooting of pennycress will be important to exploit for mobile resource capture and indirectly benefit agricultural systems by making biopores for subsequent crop roots to follow (Huang, Athmann, and Han 2020). At the seedling stage analyzed in this study no tradeoff was observed in having both a wide and deep root system with seedling vigor a promising seedling target for appropriate establishment (Atkinson et al. 2015; Pace et al. 2014). Lateral roots are important for nutrient foraging and were shown here to greatly contribute to the dicot root system width (Bao et al. 2014; Wang et al. 2002). Harnessing and matching these diverse root phenotypes to the growth environment and desired ecosystem function may be possible. Characterization of a mapping population for genetic mapping studies will allow functional genomic analysis of root traits and identify loci for crop improvement. Admixture of divergent genomes is expected to enhance climate adaptation and yield improvement (Lovell et al. 2021).

## 7 Author Contributions

MG, AEL, and CNT conceptualization; MG, AEL, SLG, NMM, and EYM experimentation; MG, AEL, and TP image processing; MG and AEL statistical analysis. All authors contributed to the writing of the manuscript and approved the final version for submission.

## 8 Acknowledgments

The authors would like to thank Tim Parker for computation assistance; Ni Jiang for DynamicRoot guidance; Ratan Chopra, John C. Sedbrook, Winthrop B. Phippen, and David Marks for providing seeds used in this study; and all iPREP project members for guidance. This study was supported by the U.S. Department of Energy, Office of Science, Office of Biological and Environmental Research, Genomic Science Program grant no. DE-SC0021286 to Christopher N. Topp.

## 10 Data Availability Statement

The dataset supporting the results of this article, the statistical analysis R codes, and modified PlantCV rosette imaging code is available online as a Zenodo repository https://doi.org/10.5281/zenodo.7536940.

