## Supplementary tables for "A temporal atlas and response to nitrate availability of 3D root system architecture in diverse pennycress (*Thlaspi arvense* L.) accessions"

### 11 Supplementary data

*Table 1. Traits measured and descriptors used in this study. Calculations for derived traits are found in the Supplementary R code.*

| Trait category | Trait description | Method | Units |
|---|---|---|---|
| Root size | Root dry mass total | Measured after 3 days at 60°C | g |
| | Total count | DynamicRoots | - |
| | Lateral count | DynamicRoots | - |
| | Total length | DynamicRoots | mm |
| | Primary length | DynamicRoots | mm |
| | Extended primary root length | Measured using a ruler | mm |
| | Lateral length | DynamicRoots | mm |
| | First order lateral length | DynamicRoots | mm |
| | Total volume | DynamicRoots | $mm^3$ |
| | Lateral volume | DynamicRoots | $mm^3$ |
| | First order lateral volume | DynamicRoots | mm |
| Root distribution | Convex hull volume of root system | GiA3D | $mm^3$ |
| | Solidity of root system | GiA3D | ratio |
| | Bushiness of root system | GiA3D | ratio |
| | Density first order laterals | DynamicRoots | TL |
| | Density first order laterals | DynamicRoots | BRTL |
| | Average interbranch distance | DynamicRoots | mm |
| | Maximum width of root system | GiA3D | mm |
| | Maximum depth of root system | GiA3D | mm |
| | Width:Depth ratio of root system | GiA3D | ratio |
| | Depth of all roots | DynamicRoots | mm |
| | Depth of all lateral roots | DynamicRoots | mm |
| | Length distribution of root system | Gia3D | mm |
| | Average tortuosity of all roots | DynamicRoots | ratio |
| | Average tortuosity of primary root | DynamicRoots | ratio |
| | Average tortuosity of lateral roots | DynamicRoots | ratio |
| | Average soil angle of all roots | DynamicRoots | degrees |
| | Primary root soil angle | DynamicRoots | degrees |
| | Average soil angle of lateral roots | DynamicRoots | degrees |
| | Average branching angle from parent | DynamicRoots | degrees |
| | Average branching angle of laterals | DynamicRoots | degrees |
| | Specific root length | Root length / Root dry mass (derived) | $m\ g^{-1}$ |
| | Root lateral:axial fraction | Lateral + secondary lateral root length / total root length (derived) | - |
| Root Growth | Lateral root growth rate | Change in lateral root length / hours (derived) | $mm\ hour^{-1}$ |
| | Total root growth rate | Change in total root length / hours (derived) | $mm\ hour^{-1}$ |
| Root anatomy | Root maximum diameter | | mm |
| | Average radius of all roots | DynamicRoots | mm |
| | Radius of primary root | DynamicRoots | mm |
| | Average radius lateral roots | DynamicRoots | mm |
| Shoot size | Shoot dry mass total | Measured after 3 days at 60°C | g |
| | Rosette size | PlantCV | cm |
| | Leaf count | PlantCV | cm |

| | | | |
|---|---|---|---|
| Biomass distribution | Plant dry mass total | Root + Shoot dry mass (derived) | g |
| | Root mass fraction | Root dry mass / total dry mass (derived) | $g\ g^{1}$ |

*Table 2. Day x N condition ANOVA for plant traits in Spring32 in four nitrate conditions*

| Trait | Source of variation | | |
|---|---|---|---|
| | Day | NTreatment | Day:NTreatment |
| InterbranchMeanDistance_mm | 95.35 *** | 5.88 ** | 2.46 * |
| InterbranchMedianDistance_mm | 170.64 *** | 3.64 * | 1.84 ns |
| RootBushiness3D | 40.14 *** | 1.26 ns | 1.68 ns |
| RootConvexHullVolume3D_mm3 | 91.91 *** | 7.39 *** | 3.58 *** |
| RootCountFirstOrderLateral | 211.66 *** | 2.72 ns | 0.80 ns |
| RootCountLateral | 93.25 *** | 4.34 ** | 2.63 * |
| RootDensityFirstOrderLateral_BRTL | 167.02 *** | 4.48 ** | 1.22 ns |
| RootDensityFirstOrderLateral_TL | 114.65 *** | 5.09 ** | 0.91 ns |
| RootDepth3D_mm | 100.41 *** | 3.91 * | 0.41 ns |
| RootDepthPrimary_mm | 65.25 *** | 3.04 * | 0.45 ns |
| RootLateralGrowthRate_mm.h | 164.01 *** | 11.36 *** | 5.18 *** |
| RootLengthDistribution3D_mm | 1041.93 *** | 11.18 *** | 1.33 ns |
| RootLengthFirstOrderLateral_mm | 199.83 *** | 14.82 *** | 5.90 *** |
| RootLengthLateral_mm | 164.01 *** | 11.36 *** | 5.18 *** |
| RootLengthPrimary_mm | 121.67 *** | 2.95 * | 0.66 ns |
| RootLengthSecondOrderLateral_mm | 19.86 *** | 0.87 ns | 1.45 ns |
| RootMaximumNetworkWidth3D_mm | 226.28 *** | 12.98 *** | 3.64 *** |
| RootMeanBranchingAngle_degrees | 124.42 *** | 0.68 ns | 1.20 ns |
| RootMeanBranchingAngleFirstOrderLateral_degrees | 808.80 *** | 4.83 ** | 2.39 * |
| RootMeanBranchingAngleLateral_degrees | 1000.69 *** | 2.84 * | 3.29 ** |
| RootMeanDepth_mm | 19.79 *** | 1.53 ns | 1.17 ns |
| RootMeanDepthFirstOrderLateral_mm | 144.16 *** | 0.73 ns | 1.18 ns |
| RootMeanDepthLateral_mm | 107.52 *** | 0.90 ns | 1.57 ns |
| RootMeanLength_mm | 22.56 *** | 5.16 ** | 0.78 ns |
| RootMeanLengthFirstOrderLateral_mm | 284.10 *** | 18.45 *** | 6.34 *** |
| RootMeanLengthLateral_mm | 191.45 *** | 8.45 *** | 2.63 * |
| RootMeanRadius_mm | 35.03 *** | 9.77 *** | 1.60 ns |
| RootMeanRadiusFirstOrderLateral_mm | 1967.11 *** | 11.17 *** | 1.51 ns |
| RootMeanRadiusLateral_mm | 1546.97 *** | 6.86 *** | 1.16 ns |
| RootMeanSoilAngle_degrees | 95.21 *** | 2.37 ns | 0.57 ns |
| RootMeanSoilAngleFirstOrderLateral_degrees | 458.29 *** | 8.68 *** | 3.12 ** |
| RootMeanSoilAngleLateral_degrees | 308.70 *** | 6.08 *** | 1.96 ns |
| RootMeanTortuosity | 34.95 *** | 4.32 ** | 2.07 * |
| RootMeanTortuosityFirstOrderLateral | 15423.97 *** | 2.96 * | 1.12 ns |
| RootMeanTortuosityLateral | 21286.41 *** | 1.42 ns | 1.09 ns |
| RootMeanVolume_mm3 | 25.46 *** | 7.91 *** | 1.29 ns |
| RootMeanVolumeFirstOrderLateral_mm3 | 256.51 *** | 28.47 *** | 7.75 *** |
| RootMeanVolumeLateral_mm3 | 190.03 *** | 12.08 *** | 2.82 ** |
| RootMedianBranchingAngle_degrees | 138.13 *** | 0.67 ns | 1.49 ns |
| RootMedianBranchingAngleFirstOrderLateral_degrees | 1073.06 *** | 8.15 *** | 3.16 ** |
| RootMedianBranchingAngleLateral_degrees | 1377.15 *** | 8.10 *** | 3.69 *** |
| RootMedianDepth_mm | 23.26 *** | 1.67 ns | 1.04 ns |
| RootMedianDepthFirstOrderLateral_mm | 150.38 *** | 1.10 ns | 1.14 ns |
| RootMedianDepthLateral_mm | 113.86 *** | 1.20 ns | 1.48 ns |
| RootMedianLength_mm | 29.42 *** | 4.69 ** | 1.08 ns |
| RootMedianLengthFirstOrderLateral_mm | 148.55 *** | 12.41 *** | 4.30 *** |
| RootMedianLengthLateral_mm | 69.33 *** | 5.11 ** | 2.26 * |
| RootMedianRadius_mm | 47.29 *** | 11.37 *** | 1.54 ns |
| RootMedianRadiusFirstOrderLateral_mm | 2325.58 *** | 16.29 *** | 2.06 * |
| RootMedianRadiusLateral_mm | 2221.19 *** | 13.63 *** | 1.82 ns |
| RootMedianSoilAngle_degrees | 99.72 *** | 2.02 ns | 0.63 ns |
| RootMedianSoilAngleFirstOrderLateral_degrees | 881.29 *** | 19.39 *** | 5.49 *** |

| | | | |
|---|---|---|---|
| RootMedianSoilAngleLateral_degrees | 486.69 *** | 7.53 *** | 2.38 * |
| RootMedianTortuosity | 30.32 *** | 0.08 ns | 2.57 * |
| RootMedianTortuosityFirstOrderLateral | 24962.16 *** | 8.38 *** | 3.13 ** |
| RootMedianTortuosityLateral | 17496.68 *** | 1.16 ns | 1.28 ns |
| RootMedianVolume_mm3 | 53.49 *** | 5.42 ** | 1.79 ns |
| RootMedianVolumeFirstOrderLateral_mm | 116.82 *** | 14.35 *** | 4.58 *** |
| RootMedianVolumeLateral_mm3 | 63.01 *** | 3.58 * | 2.07 * |
| RootRadiusPrimary | 2.88 * | 31.56 *** | 1.15 ns |
| RootSoilAnglePrimary_degrees | 12.91 *** | 9.09 *** | 1.54 ns |
| RootSolidity3D | 334.40 *** | 1.82 ns | 1.80 ns |
| RootTortuosityPrimary | 0.91 ns | 0.85 ns | 0.92 ns |
| RootTotalCount | 98.25 *** | 4.78 ** | 2.58 * |
| RootTotalDepth_mm | 88.75 *** | 1.50 ns | 1.53 ns |
| RootTotalDepthFirstOrderLateral_mm | 92.81 *** | 1.23 ns | 1.14 ns |
| RootTotalDepthLateral_mm | 76.52 *** | 1.39 ns | 1.56 ns |
| RootTotalGrowthRate_mm.h | 211.79 *** | 11.81 *** | 5.19 *** |
| RootTotalLength_mm | 209.60 *** | 11.88 *** | 5.17 *** |
| RootTotalVolume_mm3 | 191.80 *** | 21.29 *** | 5.50 *** |
| RootVolumeFirstOrderLateral_mm3 | 201.32 *** | 26.84 *** | 9.01 *** |
| RootVolumeLateral_mm3 | 145.47 *** | 16.43 *** | 6.12 *** |
| RootVolumePrimary_mm3 | 120.26 *** | 11.76 *** | 0.43 ns |
| RootWidthDepth3D_Ratio | 56.15 *** | 5.48 ** | 1.71 ns |

*** $P < 0.001$; ** $P < 0.01$; * $P < 0.05$; ns not significant

*Table 3. Genotype x N condition ANOVA for plant traits at day 17 in Spring32, MN106, and ISU89, in high N and trace N conditions*

| Trait | Source of variation | | |
|---|---|---|---|
| | Geno | NTreatment | Geno:NTreatment |
| InterbranchMeanDistance_mm | 0.84 ns | 5.92 * | 0.59 ns |
| InterbranchMedianDistance_mm | 2.83 ns | 1.29 ns | 0.63 ns |
| RootBushiness3D | 1.82 ns | 2.24 ns | 0.12 ns |
| RootConvexHullVolume3D_mm3 | 12.23 *** | 13.35 ** | 3.31 ns |
| RootCountFirstOrderLateral | 0.92 ns | 0.07 ns | 2.70 ns |
| RootCountLateral | 2.17 ns | 1.88 ns | 0.64 ns |
| RootDensityFirstOrderLateral_BRTL | 1.41 ns | 5.95 * | 1.08 ns |
| RootDensityFirstOrderLateral_TL | 0.81 ns | 0.06 ns | 1.65 ns |
| RootDepth3D_mm | 2.31 ns | 13.44 ** | 0.48 ns |
| RootDepthPrimary_mm | 1.80 ns | 13.11 ** | 0.45 ns |
| RootDW_g | 0.91 ns | 0.22 ns | 1.38 ns |
| RootLateralGrowthRate_mm.h | 2.52 ns | 21.16 *** | 0.07 ns |
| RootLengthDistribution3D_mm | 1.49 ns | 2.54 ns | 1.83 ns |
| RootLengthFirstOrderLateral_mm | 0.89 ns | 4.10 ns | 1.09 ns |
| RootLengthLateral_mm | 0.25 ns | 1.27 ns | 0.21 ns |
| RootLengthPrimary_mm | 0.65 ns | 4.69 * | 0.60 ns |
| RootLengthSecondOrderLateral_mm | 0.96 ns | 0.00 ns | 1.20 ns |
| RootMaximumNetworkWidth3D_mm | 19.82 *** | 28.85 *** | 4.17 * |
| RootMeanBranchingAngle_degrees | 6.75 ** | 0.26 ns | 0.01 ns |
| RootMeanBranchingAngleFirstOrderLateral_degrees | 11.54 *** | 1.53 ns | 0.53 ns |
| RootMeanBranchingAngleLateral_degrees | 6.86 ** | 0.32 ns | 0.01 ns |
| RootMeanDepth_mm | 0.37 ns | 0.24 ns | 1.00 ns |
| RootMeanDepthFirstOrderLateral_mm | 0.82 ns | 0.01 ns | 1.28 ns |
| RootMeanDepthLateral_mm | 0.34 ns | 0.05 ns | 1.14 ns |
| RootMeanLength_mm | 22.53 *** | 2.00 ns | 1.19 ns |
| RootMeanLengthFirstOrderLateral_mm | 12.06 *** | 9.96 ** | 1.38 ns |
| RootMeanLengthLateral_mm | 18.89 *** | 3.06 ns | 1.81 ns |
| RootMeanRadius_mm | 0.36 ns | 2.14 ns | 0.73 ns |
| RootMeanRadiusFirstOrderLateral_mm | 0.20 ns | 1.64 ns | 0.40 ns |
| RootMeanRadiusLateral_mm | 0.34 ns | 1.95 ns | 0.87 ns |
| RootMeanSoilAngle_degrees | 7.74 ** | 10.83 ** | 0.23 ns |
| RootMeanSoilAngleFirstOrderLateral_degrees | 34.86 *** | 16.52 *** | 0.40 ns |
| RootMeanSoilAngleLateral_degrees | 9.02 ** | 11.73 ** | 0.26 ns |
| RootMeanTortuosity | 0.89 ns | 0.00 ns | 0.90 ns |
| RootMeanTortuosityFirstOrderLateral | 7.44 ** | 0.13 ns | 1.62 ns |
| RootMeanTortuosityLateral | 3.21 ns | 0.89 ns | 0.79 ns |
| RootMeanVolume_mm3 | 12.82 *** | 0.04 ns | 0.21 ns |
| RootMeanVolumeFirstOrderLateral_mm3 | 13.08 *** | 8.58 ** | 0.78 ns |
| RootMeanVolumeLateral_mm3 | 17.34 *** | 2.11 ns | 0.85 ns |
| RootMedianBranchingAngle_degrees | 9.99 *** | 0.00 ns | 0.06 ns |
| RootMedianBranchingAngleFirstOrderLateral_degrees | 10.99 *** | 1.25 ns | 0.23 ns |
| RootMedianBranchingAngleLateral_degrees | 9.57 *** | 0.01 ns | 0.05 ns |
| RootMedianDepth_mm | 0.13 ns | 0.05 ns | 0.21 ns |
| RootMedianDepthFirstOrderLateral_mm | 0.42 ns | 0.60 ns | 0.56 ns |
| RootMedianDepthLateral_mm | 0.14 ns | 0.03 ns | 0.19 ns |
| RootMedianLength_mm | 9.06 ** | 0.36 ns | 2.01 ns |
| RootMedianLengthFirstOrderLateral_mm | 9.37 ** | 7.81 * | 1.95 ns |
| RootMedianLengthLateral_mm | 9.05 ** | 0.10 ns | 2.27 ns |
| RootMedianRadius_mm | 0.81 ns | 2.55 ns | 0.32 ns |
| RootMedianRadiusFirstOrderLateral_mm | 0.91 ns | 2.78 ns | 0.09 ns |
| RootMedianRadiusLateral_mm | 0.49 ns | 2.86 ns | 0.69 ns |
| RootMedianSoilAngle_degrees | 12.53 *** | 9.95 ** | 0.89 ns |
| RootMedianSoilAngleFirstOrderLateral_degrees | 32.04 *** | 13.41 ** | 0.10 ns |
| RootMedianSoilAngleLateral_degrees | 11.06 *** | 10.52 ** | 0.91 ns |
| RootMedianTortuosity | 3.59 * | 2.05 ns | 0.18 ns |
| RootMedianTortuosityFirstOrderLateral | 7.24 ** | 0.34 ns | 0.10 ns |
| RootMedianTortuosityLateral | 3.72 * | 2.82 ns | 0.09 ns |

| | | | |
|---|---|---|---|
| RootMedianVolume_mm3 | 8.58 ** | 0.51 ns | 0.59 ns |
| RootMedianVolumeFirstOrderLateral_mm | 7.79 ** | 4.53 * | 0.25 ns |
| RootMedianVolumeLateral_mm3 | 9.06 ** | 0.16 ns | 0.94 ns |
| RootRadiusPrimary | 1.38 ns | 0.09 ns | 0.32 ns |
| RootSoilAnglePrimary_degrees | 1.08 ns | 6.85 * | 0.76 ns |
| RootSolidity3D | 20.98 *** | 6.81 * | 0.09 ns |
| RootTortuosityPrimary | 1.03 ns | 2.29 ns | 0.78 ns |
| RootTotalCount | 1.93 ns | 2.24 ns | 0.57 ns |
| RootTotalDepth_mm | 1.78 ns | 0.38 ns | 1.45 ns |
| RootTotalDepthFirstOrderLateral_mm | 1.35 ns | 0.09 ns | 1.88 ns |
| RootTotalDepthLateral_mm | 1.59 ns | 0.34 ns | 1.35 ns |
| RootTotalGrowthRate_mm.h | 1.12 ns | 4.78 * | 1.10 ns |
| RootTotalLength_mm | 1.24 ns | 4.40 * | 0.64 ns |
| RootTotalVolume_mm3 | 2.32 ns | 2.82 ns | 1.56 ns |
| RootVolumeFirstOrderLateral_mm3 | 1.49 ns | 4.78 * | 0.77 ns |
| RootVolumeLateral_mm3 | 1.03 ns | 0.82 ns | 0.23 ns |
| RootVolumePrimary_mm3 | 20.87 *** | 43.39 *** | 5.98 ** |
| RootWidthDepth3D_Ratio | 1.14 ns | 0.41 ns | 0.13 ns |
| ShootDW_g | 1.77 ns | 1.15 ns | 1.74 ns |
| SpecRootLength_m.g | 0.86 ns | 0.49 ns | 0.22 ns |
| TotalMass_g | 0.84 ns | 5.92 * | 0.59 ns |

*** P < 0.001; ** P < 0.01; * P < 0.05; ns not significant

12 *Table 4. Day x Genotype ANOVA for plant traits across the 24 diverse pennycress accession lines [missing shoot traits]*

| Trait | Source of variation | | |
|---|---|---|---|
| | Day | Geno | Geno:Day |
| Extended_Root | 18.38 *** | 0.90 ns | 0.84 ns |
| InterbranchMeanDistance_mm | 96.33 *** | 2.01 ** | 1.05 ns |
| InterbranchMedianDistance_mm | 69.85 *** | 1.32 ns | 1.02 ns |
| RootBushiness3D | 107.98 *** | 4.72 *** | 1.57 ** |
| RootConvexHullVolume3D_mm3 | 164.11 *** | 7.72 *** | 2.64 *** |
| RootCountFirstOrderLateral | 243.43 *** | 3.60 *** | 1.22 ns |
| RootCountLateral | 343.20 *** | 4.57 *** | 1.20 ns |
| RootDensityFirstOrderLateral_BRTL | 78.63 *** | 1.74 * | 1.13 ns |
| RootDensityFirstOrderLateral_TL | 126.18 *** | 3.07 *** | 1.42 * |
| RootDepth3D_mm | 448.05 *** | 2.82 *** | 0.57 ns |
| RootDepthPrimary_mm | 419.96 *** | 2.70 *** | 0.57 ns |
| RootLateralGrowthRate_mm.h | 91.04 *** | 2.71 *** | 1.05 ns |
| RootLengthDistribution3D_mm | 223.94 *** | 2.15 ** | 1.59 ** |
| RootLengthFirstOrderLateral_mm | 156.78 *** | 3.90 *** | 1.07 ns |
| RootLengthLateral_mm | 179.60 *** | 4.41 *** | 1.10 ns |
| RootLengthPrimary_mm | 459.43 *** | 2.79 *** | 0.52 ns |
| RootLengthSecondOrderLateral_mm | 7.39 *** | 1.11 ns | 0.89 ns |
| RootMaximumNetworkWidth3D_mm | 210.88 *** | 9.12 *** | 1.70 ** |
| RootMeanBranchingAngle_degrees | 222.42 *** | 1.43 ns | 1.52 ** |
| RootMeanBranchingAngleFirstOrderLateral_degrees | 137.35 *** | 1.20 ns | 1.42 * |
| RootMeanBranchingAngleLateral_degrees | 169.08 *** | 1.27 ns | 1.65 ** |
| RootMeanDepth_mm | 58.11 *** | 2.16 ** | 1.35 * |
| RootMeanDepthFirstOrderLateral_mm | 288.52 *** | 3.23 *** | 1.03 ns |
| RootMeanDepthLateral_mm | 280.66 *** | 2.78 *** | 0.97 ns |
| RootMeanLength_mm | 11.99 *** | 2.43 *** | 1.87 *** |
| RootMeanLengthFirstOrderLateral_mm | 156.16 *** | 4.42 *** | 1.21 ns |
| RootMeanLengthLateral_mm | 185.57 *** | 5.48 *** | 1.46 * |
| RootMeanRadius_mm | 159.34 *** | 1.72 * | 1.16 ns |
| RootMeanRadiusFirstOrderLateral_mm | 104.08 *** | 1.09 ns | 1.24 ns |
| RootMeanRadiusLateral_mm | 124.20 *** | 1.29 ns | 1.44 * |
| RootMeanSoilAngle_degrees | 170.13 *** | 1.81 * | 1.69 ** |
| RootMeanSoilAngleFirstOrderLateral_degrees | 387.11 *** | 3.34 *** | 0.99 ns |
| RootMeanSoilAngleLateral_degrees | 468.39 *** | 4.27 *** | 1.07 ns |
| RootMeanTortuosity | 12.40 *** | 2.47 *** | 1.17 ns |
| RootMeanTortuosityFirstOrderLateral | 395.47 *** | 1.70 * | 1.43 * |
| RootMeanTortuosityLateral | 552.82 *** | 2.58 *** | 1.81 *** |
| RootMeanVolume_mm3 | 48.40 *** | 1.76 * | 1.71 *** |
| RootMeanVolumeFirstOrderLateral_mm3 | 76.05 *** | 3.27 *** | 0.86 ns |
| RootMeanVolumeLateral_mm3 | 87.91 *** | 3.90 *** | 0.92 ns |
| RootMedianBranchingAngle_degrees | 216.96 *** | 1.44 ns | 1.58 ** |
| RootMedianBranchingAngleFirstOrderLateral_degrees | 133.94 *** | 1.17 ns | 1.50 ** |
| RootMedianBranchingAngleLateral_degrees | 169.25 *** | 1.28 ns | 1.81 *** |
| RootMedianDepth_mm | 24.90 *** | 2.31 *** | 1.21 ns |
| RootMedianDepthFirstOrderLateral_mm | 243.42 *** | 3.12 *** | 1.05 ns |
| RootMedianDepthLateral_mm | 233.29 *** | 2.65 *** | 0.99 ns |
| RootMedianLength_mm | 38.94 *** | 2.14 ** | 1.78 *** |
| RootMedianLengthFirstOrderLateral_mm | 124.50 *** | 3.88 *** | 1.19 ns |
| RootMedianLengthLateral_mm | 148.14 *** | 4.70 *** | 1.54 ** |
| RootMedianRadius_mm | 219.46 *** | 1.61 * | 1.42 * |
| RootMedianRadiusFirstOrderLateral_mm | 104.24 *** | 1.10 ns | 1.30 ns |
| RootMedianRadiusLateral_mm | 124.06 *** | 1.33 ns | 1.52 ** |
| RootMedianSoilAngle_degrees | 164.46 *** | 1.83 * | 1.75 *** |
| RootMedianSoilAngleFirstOrderLateral_degrees | 390.10 *** | 3.00 *** | 1.00 ns |
| RootMedianSoilAngleLateral_degrees | 488.34 *** | 4.18 *** | 1.12 ns |
| RootMedianTortuosity | 17.34 *** | 1.63 * | 1.43 * |
| RootMedianTortuosityFirstOrderLateral | 398.32 *** | 1.75 * | 1.44 * |
| RootMedianTortuosityLateral | 559.47 *** | 2.61 *** | 1.79 *** |
| RootMedianVolume_mm3 | 108.72 *** | 1.72 * | 1.86 *** |

| | | | |
|---|---|---|---|
| RootMedianVolumeFirstOrderLateral_mm | 64.10 *** | 3.10 *** | 0.92 ns |
| RootMedianVolumeLateral_mm3 | 84.65 *** | 4.13 *** | 1.11 ns |
| RootRadiusPrimary | 2.60 ns | 2.39 *** | 0.76 ns |
| RootSoilAnglePrimary_degrees | 34.44 *** | 1.26 ns | 1.50 ** |
| RootSolidity3D | 186.13 *** | 1.88 ** | 0.91 ns |
| RootTortuosityPrimary | 1.84 ns | 2.39 *** | 0.76 ns |
| RootTotalCount | 353.14 *** | 4.67 *** | 1.09 ns |
| RootTotalDepth_mm | 216.47 *** | 2.89 *** | 0.89 ns |
| RootTotalDepthFirstOrderLateral_mm | 146.80 *** | 3.06 *** | 1.01 ns |
| RootTotalDepthLateral_mm | 160.05 *** | 2.88 *** | 0.93 ns |
| RootTotalGrowthRate_mm.h | 94.02 *** | 2.60 *** | 1.08 ns |
| RootTotalLength_mm | 361.66 *** | 5.26 *** | 1.10 ns |
| RootTotalVolume_mm3 | 211.07 *** | 3.95 *** | 0.62 ns |
| RootVolumeFirstOrderLateral_mm3 | 112.09 *** | 4.14 *** | 1.03 ns |

*** $P < 0.001$; ** $P < 0.01$; * $P < 0.05$; ns not significant
